# Sustained attention is more closely related to long-term memory than to attentional control

**DOI:** 10.1101/2025.03.13.643171

**Authors:** Chong Zhao, Anna Corriveau, Jin Ke, Edward K. Vogel, Monica D. Rosenberg

## Abstract

Individuals differ in their ability to sustain attention. However, whether differences in sustained attention reflect differences in processes related to attentional control and working memory or long-term memory (LTM) remains underexplored. In Experiment 1, we conducted an online study (n = 136) measuring participants’ sustained attention, attention control and working memory, and LTM. We measured sustained attention with an audio-visual continuous performance task (avCPT) in which participants responded to images while inhibiting responses to infrequent targets; attention control and working memory with Flanker, change localization, and Simon tasks; and LTM with recognition and source memory tests. Factor analyses revealed that sustained attention formed a distinct factor from attention control and working memory and LTM. Individual differences in the sustained attention factor robustly predicted individual differences in LTM and, to a lesser extent, attention control and working memory. In Experiment 2, to test how neural signatures of sustained attention related to attention control and working memory and LTM, we analyzed fMRI functional connectivity patterns collected as 20 participants performed the avCPT. A pre-trained connectome-based model of sustained attention predicted participants’ performance on out-of-scanner LTM, but not attention control and working memory, tasks. Together these results suggest that individual differences in sustained attention, although correlated with attention control and working memory, are more closely related to LTM.

## Introduction

“Everyone knows what attention is.” Over a century ago, William James (1890) described the concept of attention in *The Principles of Psychology*. He defined attention as the ability to concentrate on targets and effectively inhibit distractors, which he saw as essential for higher-level processes such as memory and learning (James, 1890). Importantly, James also emphasized that voluntary attention couldn’t be sustained for “more than a few seconds at a time.” Instead, he saw the goal of sustaining attention as successively bringing relevant items back into mind. Mirroring these descriptions of attention by James (1890), contemporary research divides visual attention into multiple subcomponents including sustained attention and attentional control (Chun et al., 2011; M. D. Rosenberg & Chun, 2020).

One line of attentional research has focused on participants’ ability to focus on task goals throughout a longer timeframe, or sustain attention (Esterman & Rothlein, 2019). Sustained attention was found to fluctuate across various time scales (M. D. Rosenberg et al., 2020). Sustained attentional function is typically measured with continuous-performance tasks (CPTs), in which participants are instructed to respond to frequent targets and inhibit response to infrequent non-targets (Robertson et al., 1997; Rosenberg et al., 2013). In assessing sustained attention, errors in responding to infrequent non-targets are considered lapses in attention, while greater variability in response times is indicative of lower attentional states (Chidharom et al., 2021; Esterman et al., 2013; Rosenberg et al., 2013).

In studying another aspect of attention, attentional control, researchers have primarily focused on how individuals prioritize task-relevant information, effectively attending to targets and filtering out distractors (Burgoyne et al., 2023). Tasks measuring attentional control abilities often require participants to resolve conflicting cognitive and motor demands (Draheim et al., 2022) or detect targets amidst multiple distractors (Martin et al., 2021). Researchers have argued that this ability impacts complex cognitive task performance (Burgoyne & Engle, 2020). The ability to control attention is closely related to the function of working memory, a short-term memory store that deals with the maintenance of targets and the filtering of distractors (Vogel et al., 2005). Suggesting that these two processes share common neural mechanisms, a multivariate classifier trained on fMRI data to predict attentional selection can also decode working memory content (Zhou et al., 2022). Crucially, individual differences in working memory capacity have been shown to predict performance on attentional control tasks, including the antisaccade task, Stroop task, and Flanker task (Colom et al., 2008; Kane et al., 2001; Meier & Kane, 2017; Schweizer & Moosbrugger, 2004; Unsworth et al., 2014). Given the strong correlations among these measures, we combined them into a single attentional control and working memory factor for our analyses, while still reporting correlations for each individual working memory and attentional control task.

Because sustained attention and attentional control and working memory are typically studied separately, an open question is how they are related to each other. One possibility is that variance in sustained attention and attentional control is highly overlapping. Supporting this perspective, one study measured performance on multiple attention tasks, including complex working memory and attentional control tasks, and found evidence for a general attention factor, though it did not test sustained attention (Huang et al., 2012). In addition, individual differences in both sustained attention (Corriveau et al., 2025) and attentional control (Zhao & Vogel, 2024) predict long-term memory differences, suggesting that both forms of attention play significant roles in the formation and maintenance of long-term memory. Though preliminary behavioral evidence suggested that these two attentional processes may share overlapping variance, few neural studies have been done to explore the relationship between sustained attention and attentional control.

Recent advances in neuroimaging methods suggest that common functional brain networks underlie different forms of attention. A growing body of work uses individual differences in patterns of fMRI functional brain connectivity—coordinated activity between spatially distinct brain regions—to predict individual differences in task performance and behavior (Finn et al., 2015; Shen et al., 2017). This work has revealed, for example, that functional connectivity patterns reliably predict abilities such as sustained attention in novel individuals and datasets (Rosenberg et al., 2016). The same model that predicts individual differences in sustained attention task performance also predicted overall performance on the Attention Network Task, which assesses alerting, orienting, and executive control (Rosenberg et al., 2018). Furthermore, a similar connectome-based model trained to predict performance on a sustained attentional task (i.e., CPT) predicted performance on attentional control and working memory tasks (Avery et al., 2020).

An alternative perspective is that sustained attention and attentional control, though related, are distinct types of attention. Specifically, sustained attention affects the monitoring of ongoing cognitive processes, while attentional control is more closely related to task switching and working memory capacity (Chun et al., 2011). Empirical evidence supporting this claim has shown that lapses in attention, measured through in a psychometric vigilance task requiring participants to respond to changes in color on the screen for 5-10 minutes, sustained attention to cues, and whole report working memory tasks form a distinct factor from attentional control measured with Stroop and visual search tasks (Unsworth et al., 2021). Crucially, a two-factor model with distinct attentional control and lapse factors outperformed a one-factor attention model, suggesting that a separate lapse factor would fit the empirical data better than combining both forms of attention into one factor. However, whether attentional control and sustained attention play similar roles in affecting visual long-term memory differences remained unresolved. Moreover, no neural measures were collected for previous individual difference studies.

Our primary goal was to characterize individual differences in sustained attention, attention control and working memory, and long-term memory. Firstly, with an online sample (n = 136), we asked whether and how individual differences in sustained attention reflected differences in attentional control and working memory. We next tested whether sustained attention explained unique variance in long-term memory differences even when accounting for attentional control differences, given that both sustained attention (i.e., incidental recognition memory, Corriveau et al., 2025) and attention control and working memory (i.e., intentional recognition memory, Zhao & Vogel, 2024) predict long-term memory. Lastly, using fMRI data from a distinct sample of 20 individuals, we tested whether a validated connectome-based model of sustained attention predicted out-of-scanner sustained attention, attention control and working memory, and long-term memory performance long after participants were scanned (0.5-1.5 years). Collectively, this work fills the gap in understanding how sustained attention, both behaviorally (Experiment 1) and neurally (Experiment 2), is related to attentional control, working memory, and long-term memory.

## Experiment 1 (Behavioral)

In Experiment 1, we aimed to examine relationships between sustained attention, attention control, working memory, and long-term memory. Some studies have proposed that sustained attention and attention control might be explained by a single underlying factor (i.e., Huang et al., 2012), while other research indicated that these may be distinct constructs (i.e., Unsworth et al., 2021). With an online sample (n = 136), we are able to dissect the latent psychological factors underlying performance on seven attention and memory tasks through both exploratory and confirmatory factor analyses. The sample size was estimated based on a medium effect size (Zhao & Vogel, 2024, *r* = 0.3) from prior research, with a power = 0.9 (estimated N = 113).

## Methods

### Sustained Attention Task

In Experiment 1, data were collected from an online platform (Prolific). We recruited participants that were living in the United States, 18-35 years old, with normal or corrected-to-normal vision, no color blindness, no ongoing psychological and neurological disorder, and 90% or above approval rate for previous studies. 150 participants were recruited, and 136 participants performed numerically above chance for all tasks (inclusion criteria used in Burgoyne et al., 2023; Zhao et al., 2022). Study procedures were approved by the University of Chicago Institutional Review Board and participants were compensated ($15/hr) for their participation.

During the sustained attention task phase of the study, participants completed a 10-minute audio-visual continuous performance task (avCPT; Corriveau et al., 2025, see **Fig. 1G**). Throughout the avCPT, trial-unique images and sounds were presented simultaneously, with images displayed for 1 second each and sounds lasting 1 second with a 200 ms inter-trial interval to distinguish individual sounds. Each task run included 500 trials, with images of indoor and outdoor scenes sourced from the SUN image database (Xiao et al., 2010) and sound stimuli consisting of natural and manmade sounds cropped to 1 second from online databases. Full details on stimulus curation are provided in Corriveau et al. (2025). Participants were instructed to press a button for frequent stimuli (90%) from the task-relevant modality and to withhold a response for infrequent stimuli (10%). The non-relevant modality stimuli were to be ignored. All participants in Experiment 1 were assigned to the visual condition, responding to images and ignoring sounds. The frequent stimulus category was counterbalanced across participants (i.e., indoor vs outdoor). Correct responses for frequent trials required a button press before the onset of a new stimulus (within 1200 ms of trial start). Performance during the avCPT was calculated as sensitivity (*d*’), as well as RT consistency (reciprocal of the variation of response times, 1/(variance RT/mean RT)).

**Fig 1.**
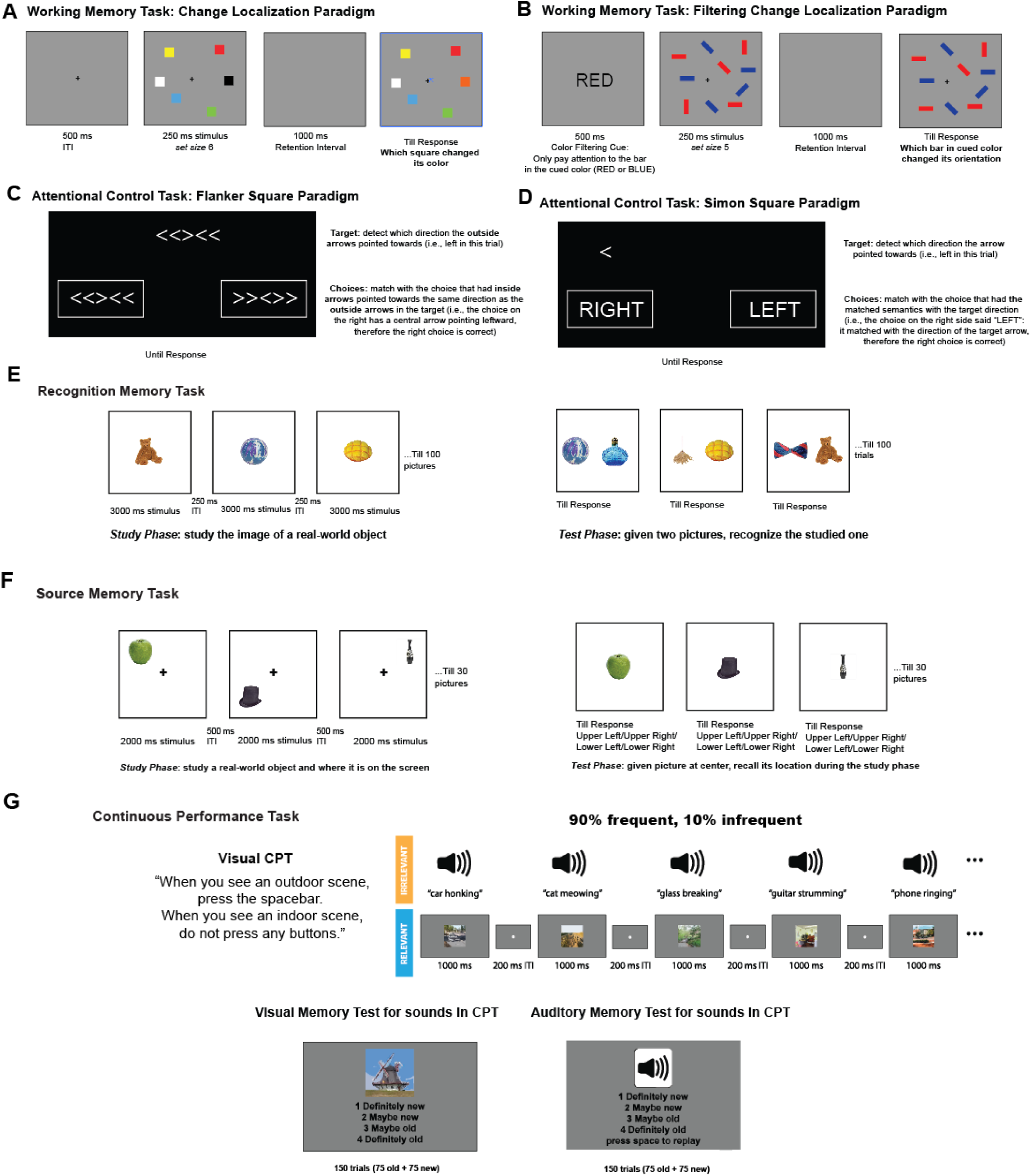
Working Memory, Attentional Control, Long-Term Memory and Sustained Attention Tasks used in Exp 1. (A) Schema of Change Localization Task. (B) Schema of the Filtering Change Localization Task. (C) Schema of the Flanker Square Task. (D) Schema of the Simon Square Task. (E) A sample of the source memory paradigm. (F) A sample of the recognition memory paradigm. (G) A sample of the av-CPT task with visual stimuli as the relevant stimuli.

#### Visual and Auditory Memory Task for avCPT stimuli

Following the avCPT, participants underwent recognition memory testing for the stimuli presented (see **Fig. 1G**). Recognition for stimuli in the task-relevant modality (images in Experiment 1) was assessed first, followed by recognition for stimuli in the task-irrelevant modality (sounds). Each memory task, for both task-relevant and irrelevant stimuli, consisted of 150 trials (75 old and 75 new). In each task, memory stimuli included 25 old stimuli from the frequent category, 25 old stimuli from the infrequent category, 25 stimuli that had been paired with the infrequent category of the opposite modality (always from the frequent category of the modality being tested), and 75 category-matched foil stimuli. Participants reported their confidence in their recognition memory on a scale from 1 to 4 (1 = definitely new, 2 = maybe new, 3 = maybe old, 4 = definitely old). The memory task was self-paced, ending each trial after a memory judgment was made. However, to prevent participants from pausing and resuming after an extended period, memory trials timed out if no response was made within 20 seconds. Timed-out trials were excluded from analyses. Images remained on the screen until a memory judgment was made. Sounds were presented at the trial’s onset, and participants were able to replay sounds as many times as needed before making a memory judgment.

### Attention Control Tasks

#### (1) Flanker Square

The Flanker Square task was adapted from the Flanker task (Burgoyne et al., 2023, see **Fig. 1C**). In each trial, participants were presented with a target stimulus and two possible response options, both consisting of sets of five arrows arranged horizontally (e.g., <<><<). Participants were instructed to choose the response option where the middle arrow aligned in direction with the outer arrows of the target stimulus. For example, if the target stimulus displayed arrows pointing left and right (e.g., <<><<), participants were to select the response option with a central arrow pointing left (e.g., >><>>). Thus, the task required participants to focus on the outer arrows of the target stimulus and the central arrow of the response options, while ignoring the central arrow of the target stimulus and the outer arrows of the response options. Each participant completed 30 seconds of practice followed by a 90-second test phase. The Flanker Square score was calculated as the difference between the number of correct and incorrect responses. Higher scores in Flanker Square corresponded to better attentional control abilities.

#### (2) Simon Square

The Simon Square task was adapted from the Simon task (Burgoyne et al., 2023, see **Fig. 1D**). In each trial, participants were presented with a target stimulus, an arrow, and two response options: the words “RIGHT” and “LEFT.” Participants were instructed to select the response option that matched the direction indicated by the arrow. For example, if the arrow pointed to the left, participants would choose the response option with the word “LEFT.” The target stimulus and response options could appear on either side of the screen with equal probability. Therefore, participants needed to focus on the direction of the arrow while interpreting the response options’ meaning, ignoring the screen’s side where the stimuli appeared. Each participant completed 30 seconds of practice followed by a 90-second test phase. The Simon Square score was calculated as the difference between the number of correct and incorrect responses. Higher scores in Simon Square corresponded to better attentional control abilities.

### Working Memory Tasks

#### (1) Change Localization

The Change Localization task was derived from the color Change Localization task utilized in previous studies (Zhao et al., 2022, see **Fig. 1A**). In each trial, six colored squares were presented simultaneously for 250 ms, followed by a blank retention interval of 1,000 ms. Subsequently, the same six squares reappeared in their original positions, with one color altered to a hue not previously presented in that trial. Each square was numbered from 1 to 6, and participants were instructed to press the corresponding key to identify the square that had changed color. The spatial arrangement of the six numbers was randomized across trials. Our behavioral measure of interest was accuracy in the Change Localization task.

#### (2) Filtering Change Localization

The Filtering Change Localization task was adapted from the Filtering Change Detection task (Luck & Vogel, 1997; Martin et al., 2021, see **Fig. 1B**). In each trial, a word, either “RED” or “BLUE,” indicating the color of the items to be attended to (the selection instruction), was presented for 200 ms, followed by a 100-ms interval. Subsequently, 10 bars were displayed for 250 ms, with half of them in the designated color, creating a set size 5 condition. After a 900-ms delay, only the bars in the attended color reappeared. During the test phase, one of the bars changed its orientation compared to the encoding phase. Participants were required to identify which one of the five bars had changed its orientation. This Filtering Localization phase comprised a total of 60 trials. Our behavioral measure of interest was the accuracy in the Filtering Change Localization task.

### Long-term Memory Tasks

#### (1) Recognition Memory Task (Behavioral) Data Acquisition

In the study phase, participants were presented with 100 images of real-world objects (see **Fig. 1E**). The average picture size was approximately 4.6° by 4.6° of visual angle, assuming a viewing distance of 80 cm from the screen. Each image was centered on the screen during both the study and test phases.

Each trial in the study phase began with a 250-ms fixation cross, followed by the display of an image at the center of the screen for 3000 ms. In the test phase, participants were presented with 100 pairs of images, one on the left side of the screen and the other on the right side. Each pair consisted of one image from the study phase and one novel image. Participants performed a two-alternative forced choice task to identify which image had been presented during the study phase.

Each test phase trial began with a 250-ms fixation cross, followed by the presentation of the test images until the participant made a button press to indicate their old versus new response and confidence level. Participants used the number keys on a keyboard to indicate their confidence and which image was previously studied: the number keys 1 and 2 for the left-side image with high and low confidence, respectively, and the number keys 9 and 8 for the right-side image with high and low confidence, respectively. The stimuli were sourced from a previously published set of real-world objects (Brady et al., 2008). Our behavioral measure of interest was the accuracy in the Recognition Memory task.

#### (2) Source Memory Task (Behavioral) Data Acquisition

During the study phase, participants were presented with 30 images of real-world objects, each averaging approximately 4.6° by 4.6° of visual angle, assuming a viewing distance of 80 cm from the screen (Zhao & Vogel, 2025, see **Fig. 1F**).

Each trial in the study phase began with a 500-ms fixation cross, followed by the display of an image at the center of the screen for 2000 ms. Participants were instructed to memorize both the visual appearance and the spatial location of each image. In the test phase, participants were presented with the same 30 images, one at a time, and were asked to press W/E/S/D to indicate which quadrant the image was originally located in (W for upper left, E for upper right, S for lower left, and D for lower right).

Each test phase trial began with a 500-ms fixation cross, followed by the presentation of the test image at the center of the screen until the participant made a button press to record their source memory response. The stimuli were drawn from a previously published set of real-world objects (Brady et al., 2008). Our behavioral measure of interest was the accuracy in the Source Memory task.

## Results

### Individual differences in sustained attention predict attentional control, working memory, and long-term memory measures

We first investigated if individual differences in sustained attention were related to attentional control, working memory, and long-term memory. Our first metric of sustained attention was avCPT *d’*. A higher *d’* suggested that participants were more likely to correctly respond to frequent targets and inhibit their responses when infrequent-category images were shown on screen. We found that participants with higher levels of sustained attention (i.e., a higher *d’*) also showed better performance in attentional control (Simon square: *r*(134) = 0.42; Flanker square: *r*(134) = 0.34), working memory (Change localization: *r*(134) = 0.39; Filter change localization: *r*(134) = 0.29, *p*s < 0.01), and long-term memory (Recognition memory: *r*(134) = 0.42; Source memory: *r*(134) = 0.48; Incidental Visual Recognition Memory of CPT: *r*(134) = 0.32, *p*s < 0.01) tasks (**Fig. 2**).

**Fig 2.**
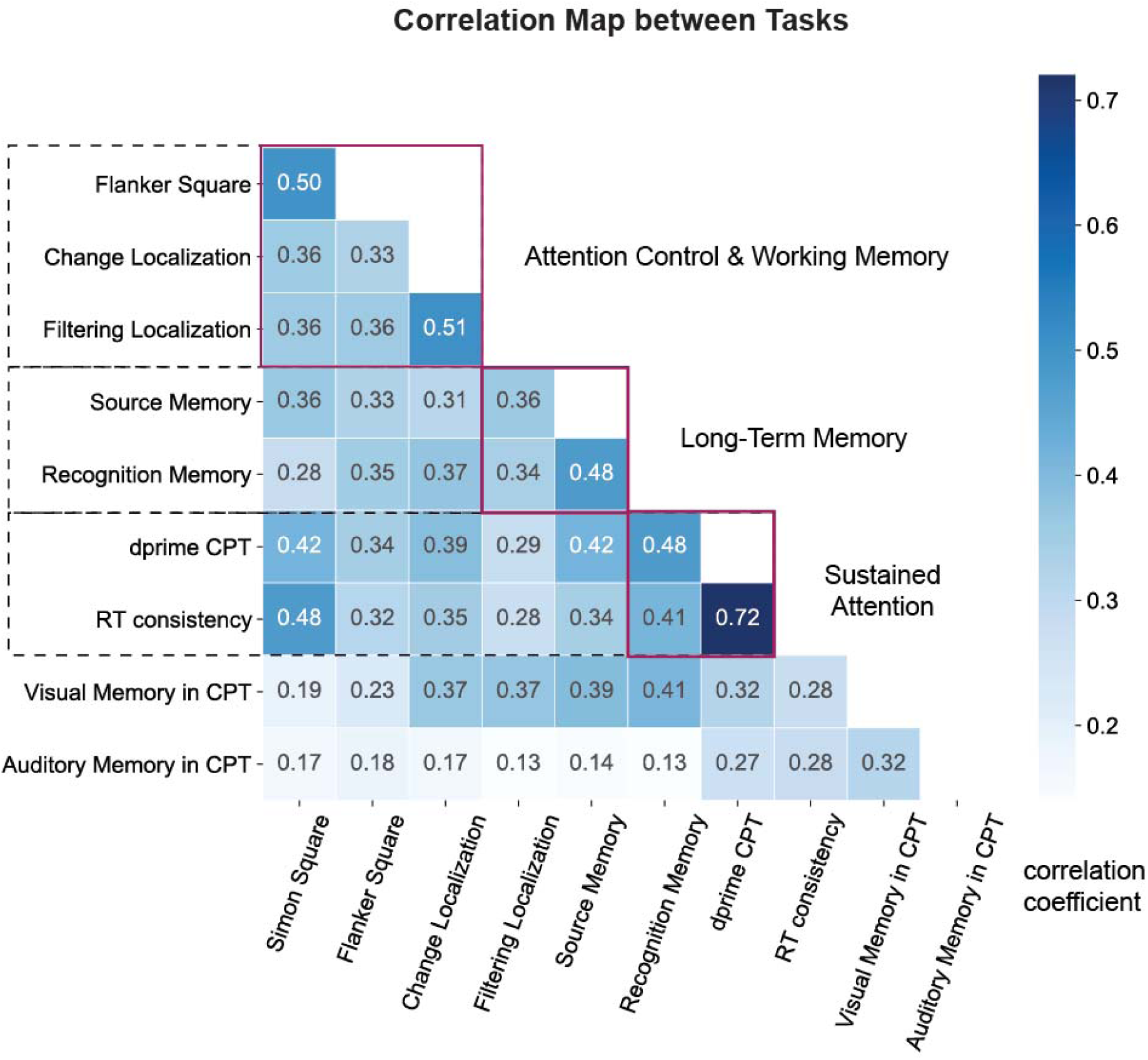
Spearman Correlation matrix of individual differences in Attentional Control Working Memory, Long-term Memory and Sustained Attention Tasks. The red squares highlighted the correlations between tasks within the same hypothesized construct (Attention Control and Working Memory, LTM and SA, respectively), which were supposed to be higher than with other tasks.

Researchers have also quantified sustained attention in terms of the fluctuations of response time during the task. The higher the variance of the response times across a session, the more attention lapses participants tend to experience (Chidharom et al., 2021; Esterman et al., 2013). Therefore, higher standardized RT variances suggested lower sustained attention levels. With standardized RT variance as our measure of sustained attention, we found that participants with higher RT consistencies (i.e., lower RT variations) showed better performance in attentional control and working memory tasks (Simon square: *r*(134) = 0.48; Flanker square: *r*(134) = 0.32; Change localization: *r*(134) = 0.35; Filter change localization: *r*(134) = 0.28, *p*s < 0.01), as well as long-term memory tasks (Recognition memory: *r*(134) = 0.34; Source memory: *r*(134) = 0.41; Incidental Visual Recognition Memory of CPT: *r*(134) = 0.28, *p*s < 0.01, see **Fig. 2**).

### Exploratory Factor Analysis suggested that a three-factor model best explains performance on attention and memory tasks

The task battery included four distinct families of cognitive measures: sustained attention, attention control, working memory, and long-term memory. To investigate how these measures clustered together, we first conducted an exploratory factor analysis on the data. Our task performance metrics were standardized (i.e., z-scored across participants) to ensure comparability across variables. Then, we used the Kaiser criterion, in combination with scree plots, to determine our optimal number of factors.

We found that the first three eigenvalues were larger than or equal to 1 (eigenvalues of 4.5, 1.1 and 1.0, respectively). Therefore, we concluded that a three-factor model best fit our task structure (18.85%, 18.82% and 14.17% variance explained respectively, 51.85% explained with all three factors). With the model specification, principal axis factoring was used for factor extraction, and an oblique promax rotation was applied to allow for correlations between factors. Finally, factor loadings were examined, with loadings greater than 0.30 deemed significant for interpreting the factors (see **Supp.** Fig. 1).

In our exploratory factor analysis, *d*’ and RT consistency in the avCPT loaded significantly onto Factor 2, suggesting that the factor reflected sustained attention abilities. Additionally, long-term memory measures, source and recognition memory, loaded positively onto Factor 1. The visual memory test for stimuli used in the avCPT and working memory tasks also loaded positively on Factor 1. These findings suggested that Factor 1 was likely a visual memory factor. Lastly, Simon Square, Flanker Square, Change Localization and Filtering Localization all positively loaded onto Factor 3, indicating that this factor reflected attentional control and working memory abilities (attention control and working memory). The auditory memory test for task-irrelevant stimuli used in the CPT did not significantly load onto the long-term memory factor. Therefore, incidental auditory recognition memory performance was excluded from further model-buildings with confirmatory factor analysis.

### Confirmatory Factor Analysis Model Comparison supported the validity of the three-factor model

In our Exploratory Factor Analysis, a three-factor model was preferred, and we assumed that the three factors reflected sustained attention (SA), Attentional Control and Working Memory, and long-term memory (LTM). To confirm that the statistical model fit with our theoretical prediction, we performed a Confirmatory Factor Analysis on our data.

The first model we proposed was a three-factor model, with sustained attention, attentional control and working memory, and long-term memory as separate factors (**Fig. 3A**). In this model, Simon Square, Flanker Square, Change Localization and Filtering Change Localization loaded onto the attention control and working memory factor. Furthermore, Source Memory, Recognition Memory and avCPT Visual Memory were loaded onto the LTM factor. Lastly, avCPT RT consistency and *d*’ were loaded onto the SA factor. The model achieved a satisfactory fit, with a CFI (Comparative Fit Index, higher = better) of 0.94, TLI (Tucker–Lewis Index, higher = better) of 0.91, and RMSEA (Root Mean Square Error of Approximation, lower = better) of 0.093. Therefore, our findings suggested that sustained attention formed a distinct factor from attentional control and long-term memory.

**Fig 3.**
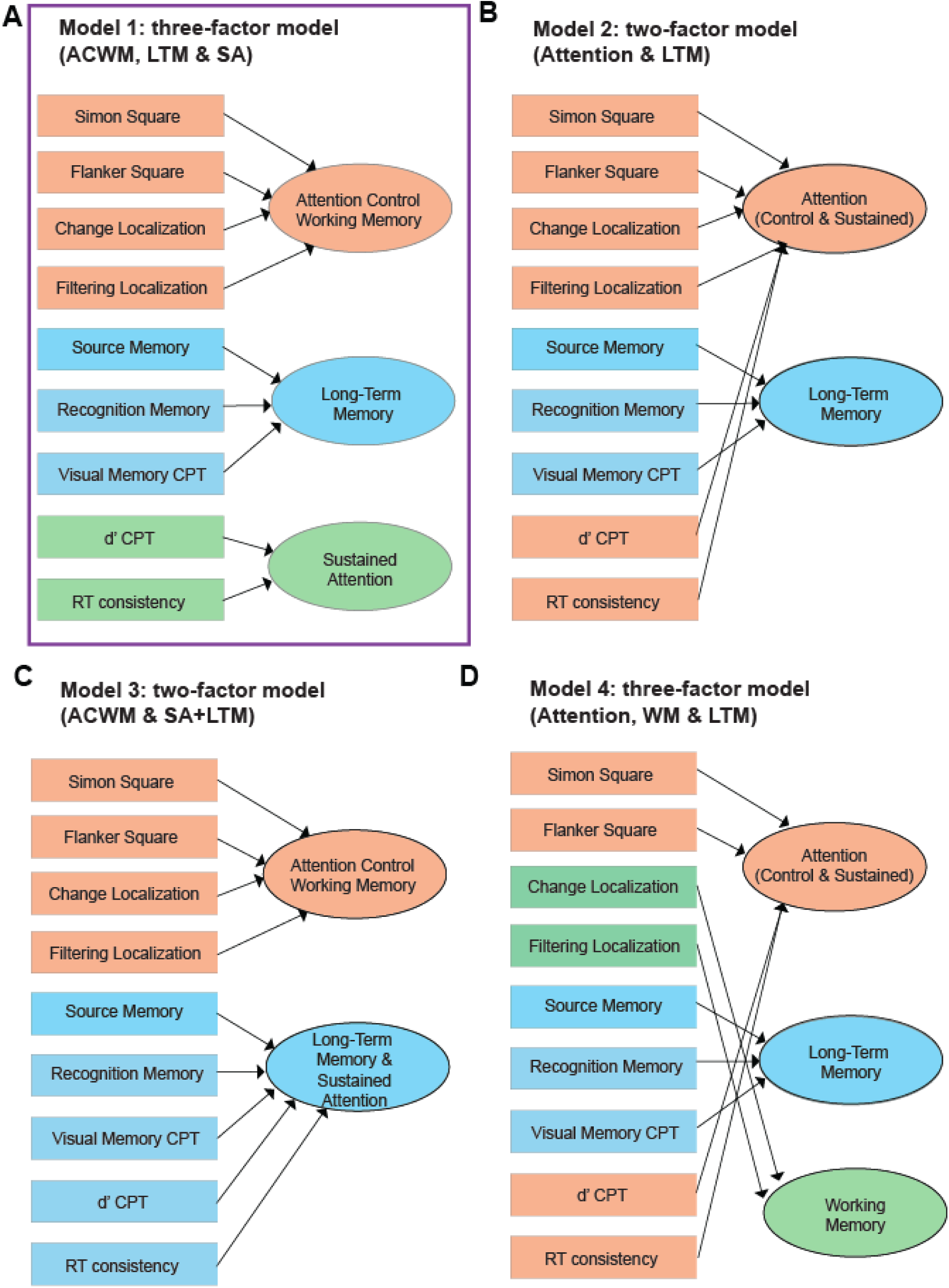
Confirmatory Factor Analysis Results. The model structures of all our potential two and three factor models. The best fitting model was highlighted in the purple box (Panel A), in which attention control and working memory, long-term memory (LTM) and sustained attention (SA) tasks loaded separately onto three distinct factors. The model fits for all four models were displayed in **Table 1**.

**Table 1.**
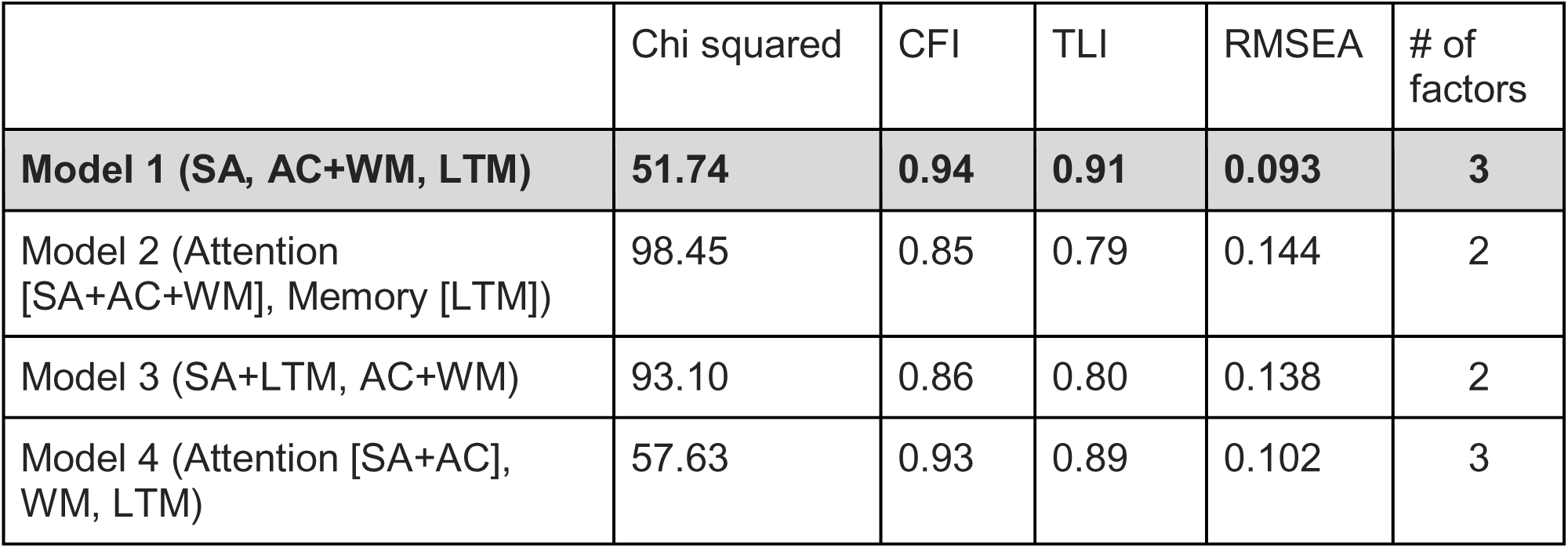
Goodness of fit for Confirmatory Factor Analysis Models. A lower Chi squared and RMSEA, and a higher CFI and TLI suggested a better model fit. Model 1 achieved the lowest Chi squared and RMSEA, and a highest CFI and TLI. Therefore, Model 1 (SA, attention control and working memory, LTM) was selected as the best model among the 4 possible CFA models.

In addition to the three-factor model that converged with our Exploratory Factor Analysis, we also examined two two-factor models that resembled the predictions of the unitary attention model. Firstly, we proposed a two-factor model that combined sustained attention, attentional control, and working memory into one attention factor, with the second factor as long-term memory (**Fig. 3B**). The model achieved reasonable fit (CFI = 0.85, TLI = 0.79, and RMSEA = 0.144; **Table 1**). Additionally, we also proposed a two-factor model that allowed sustained attention and long-term memory tasks to be our first factor, with attentional control and working memory (attention control and working memory) tasks forming a distinct second factor (**Fig. 3C**). This model achieved a similarly reasonable fit (CFI = 0.86, TLI = 0.80, and RMSEA = 0.138), slightly better than the two-factor unitary attention model, suggesting that sustained attention may be closer to long-term memory tasks than attentional control tasks. Crucially, both of the two-factor models fit worse than the three-factor model.

Lastly, we proposed an alternative three-factor model, with an attention factor (Simon Square, Flanker Square, *d’* in CPT and RT consistency), a working memory factor (Change Localization and Filtering Localization), and a long-term memory factor (Source Memory and Recognition Memory). This model achieved a similarly reasonable fit (CFI = 0.93, TLI = 0.89, and RMSEA = 0.102, **Fig. 3D**), better than the two-factor models but slightly worse than the three-factor model with sustained attention, attentional control/working memory, and long-term memory. Therefore, our findings endorsed sustained attention as a separate factor from attentional control and long-term memory (model fits shown in **Table 1**).

Under the three-factor model, we proposed that the two Square tasks and working memory tasks loaded on a Attentional Control (AC) and Working Memory (WM) Factor, long-term memory tasks loaded on a Long-Term Memory (LTM) Factor, and *d’* and RT consistency in avCPT loaded on a Sustained Attention (SA) Factor. The Confirmatory Factor Analysis loadings were all significant with our proposed framework (loadings >= 0.59, **Fig. 4A**). Moreover, the covariance values between our three factors were all significant, suggesting that AC and WM, LTM and SA were related to each other on the construct level (**Fig. 4B**). To further validate our findings, we also tried to fit a 4-factor model, with separate AC, WM, LTM, and SA factors. We found that all four factors were related to each other on the construct level, replicating our findings with the 3-factor model combining Attentional Control and Working Memory together (see **Supp.** Fig. 2).

**Fig 4.**
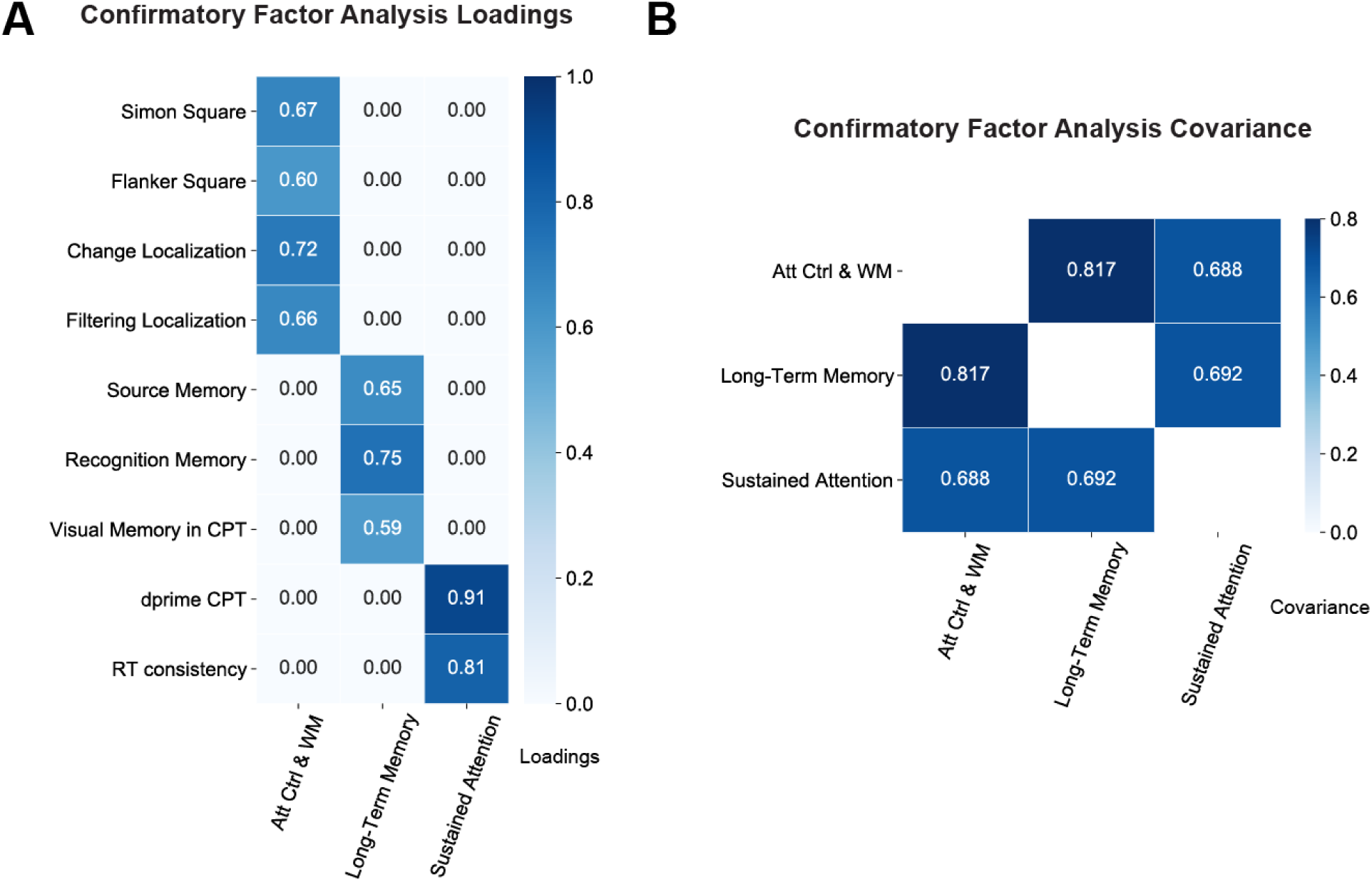
Confirmatory Factor Analysis Results. (A) The CFA model fit with Attention Control and Working Memory, LTM and SA factors shown in Fig. 4A. The loadings were all significantly positive, indicating a good model fit for our Model 1. (B) The covariance loadings between Attention Control and Working Memory, LTM and SA factors. The three factors were positively loaded onto each other.

### Individual differences in sustained attention predicted long-term memory more robustly than working memory and attentional control

With our three-factor model, we next tested whether sustained attention (SA) was more robustly related to attentional control and working memory or long-term memory (LTM). We first calculated composite scores separately for SA, attention control and working memory, and LTM. To account for potential error variances not captured by the Confirmatory Factor Analysis covariance matrix, we performed a correlational analysis relating the three factors. Results suggested that attention control and working memory and LTM were significantly correlated (*r*(134) = 0.51, *p*s < 0.01, **Fig. 5A**). More crucially, sustained attention differences significantly predicted differences in attention control and working memory (*r*(134) = 0.45, *p*s < 0.01, **Fig. 5B**) and LTM (*r*(134) = 0.73, *p*s < 0.01, **Fig. 5C**). In comparing the two correlation coefficients, we found that LTM predicted sustained attention more robustly than attention control and working memory measures predicting sustained attention (*z*(134) = 4.48, *p* < 0.01).

**Fig 5.**
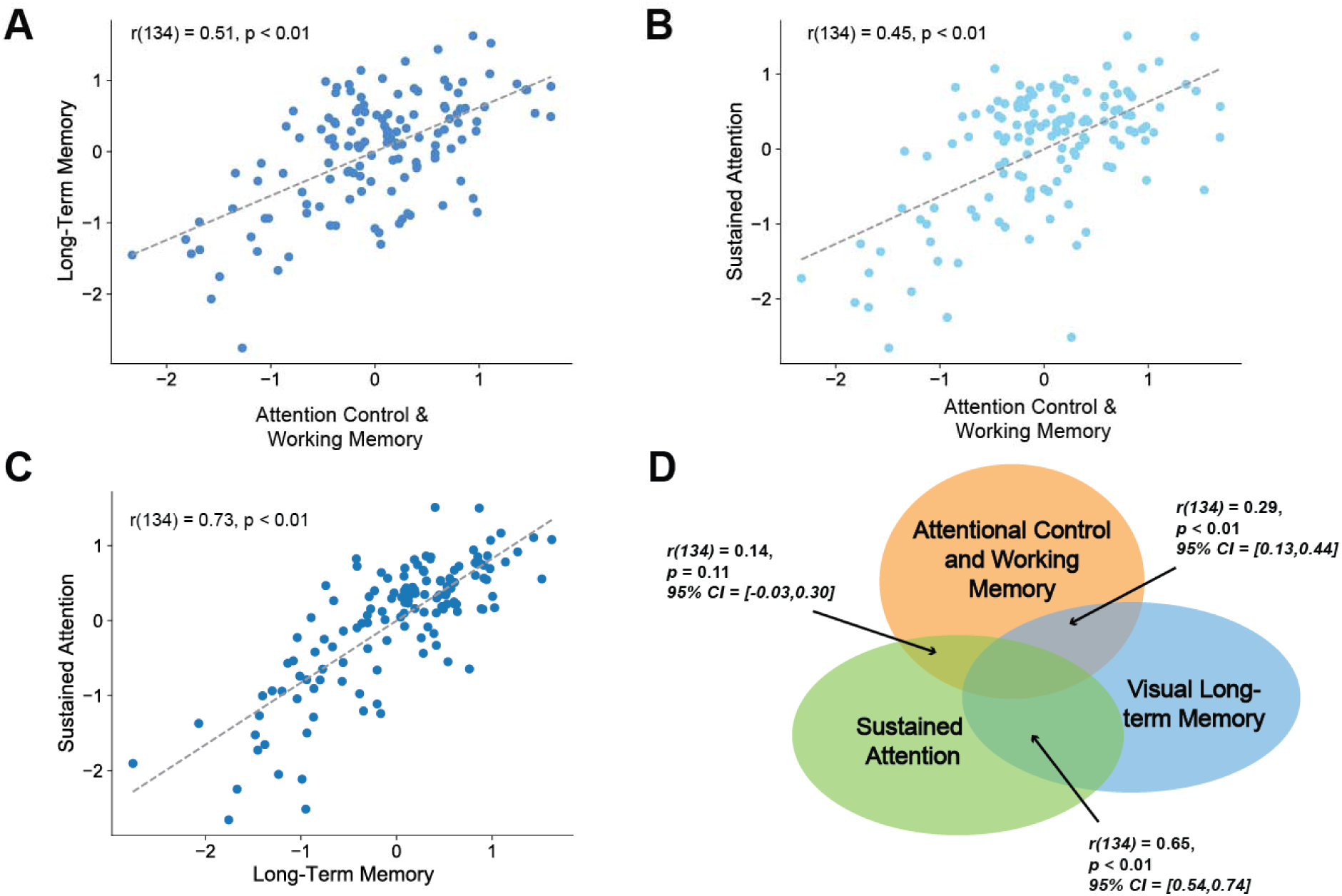
Visual Long-term Memory predicted Sustained Attention more robustly than Attentional Control and Working Memory abilities. (A) Attentional Control Working Memory Differences predicted Long-Term Memory differences. (B) Attentional Control Working Memory Differences predicted Sustained Attention differences. (C) Long-Term Memory Differences predicted Sustained Attention differences. (D) Partial regression between Attention Control and Working Memory, LTM and SA. LTM and SA shared significantly larger variance than Attention Control and Working Memory, and SA.

To control for general cognitive abilities that may affect all three factors, we ran a partial correlation analysis on SA, attention control and working memory, and LTM performance. We found that despite the significantly shared variance between each two out of the three constructs (**Fig. 6D**), LTM predicted sustained attention more robustly than attention control and working memory predicted sustained attention (*z*(134) = 5.17, *p* < 0.01). Therefore, our result converged with the Confirmatory Factor Analysis model comparison (Model 3) in that sustained attention was more closely related to long-term memory abilities than attentional control or working memory abilities.

**Fig 6.**
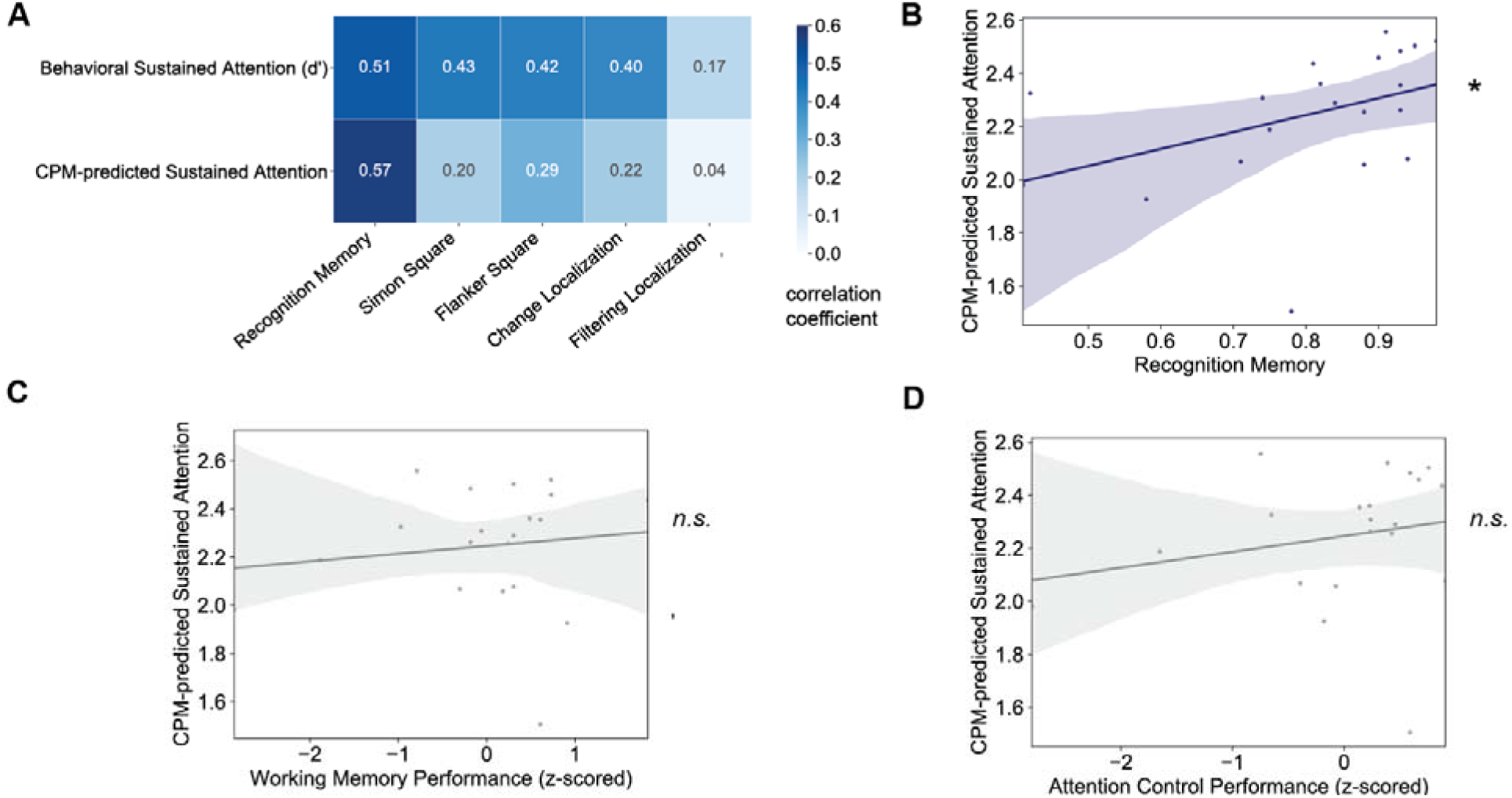
Connectome-predicted Sustained Attention predicted Recognition Memory more robustly than Attentional Control and Working Memory. (A) The correlation between behaviorally and neurally predicted sustained attention (column) and offline behavioral measures of Attention Control and Working Memory, and LTM (row). (B) Recognition Memory and saCPM-predicted sustained attention were significantly correlated. (C) Working Memory and saCPM-predicted sustained attention were not significantly correlated. (D) Attentional Control and saCPM-predicted sustained attention were not significantly correlated.

## Experiment 2 (fMRI)

In Experiment 2, we sought to conceptually replicate our findings from Experiment 1, where sustained attention was a stronger predictor of long-term memory than attentional control and working memory. Extending previous behavioral findings, we tested if neural markers of sustained attention predicted behavioral measures of attention control and working memory and LTM. In Experiment 2, we aimed to generalize our results in Experiment 1 even when the LTM and attention control and working memory tasks were administered 0.5 to 1.5 years after the sustained attention fMRI scanning session. Furthermore, we were interested in if neural markers of sustained attention, calculated with fMRI connectivity, resembled the behavioral findings that robustly predicted long-term memory.

## Method

### Participants

In Experiment 2, data were collected at the MRI Research Center at the University of Chicago. Participants (N=60) took part in at least one session of a two-session fMRI study, with sessions 10.90 days apart on average (SD = 9.61). During each session, participants completed a 10-minute audio-visual continuous performance task (avCPT; Corriveau et al., 2025), performed an annotated rest task, and watched or listened to narratives. Only avCPT data are analyzed here. Study procedures were approved by the relevant University of Chicago Institutional Review Board and participants were compensated for their participation.

### Sustained Attention Task (fMRI) Data Acquisition

Functional MRI data were collected on a 3T Philips Ingenia scanner at the MRI Research Center at the University of Chicago. Functional images were collected with a 32-channel head coil (TR/TE = 1000/28 ms, flip angle 62°, whole-brain coverage 27 slices of 3 mm thickness, in-plane resolution 2.5 × 2.5 mm2, matrix size 80 by 80, FOV 202 × 202 mm2, and MultiBand SENSE factor 3). Structural images were acquired using a high-resolution T1-weighted MPRAGE sequence (1.000 mm3 resolution, Corriveau et al., 2025). The avCPT task was the same as in Experiment 1, with the exception that images were presented for the full trial duration, 1.2 seconds, with no ITI. Performance during the avCPT was calculated as sensitivity (*d’*).

#### fMRI preprocessing

Three volumes were discarded at the beginning of each scan. The functional MRI data were preprocessed using a consistent pipeline in AFNI (Corriveau et al., 2025). Preprocessing steps included discarding initial TRs as specified, aligning the functional images to MNI space, and regressing out nuisance covariates. These covariates included a 24-parameter head motion model (6 motion parameters, their temporal derivatives, and squared terms), mean signals from subject-specific white matter and ventricle masks, and the global mean signal. Volumes were censored if the derivative of motion parameters exceeded 0.25 mm or if more than 10% of the brain was classified as outliers.

### Recognition Memory, Attention Control, and Working Memory Tasks

In Experiment 2, the recognition memory, attention control, and working memory tasks were the same as the one used in Experiment 1. We recontacted the participants 0.5-1.5 years after the fMRI scan sessions in which they performed the avCPT. 20 out of 60 participants performed the recognition memory, attention control, and working memory tasks online. Study procedures were approved by the relevant University of Chicago Institutional Review Board and participants were compensated ($20/hr) for their participation.

## Results

### Behavioral and CPM-predicted sustained attention coherently predict LTM but not attention control and working memory measures

Sustained attention (i.e., avCPT) performance during the scan session predicted performance on a recognition memory task collected 0.5-1.5 years later (*r*(18) = 0.51, *p* < 0.01, **Fig. 6A**), replicating our findings in Experiment 1 that sustained attention is closely related to long-term memory. The relationship between CPT performance and attention control and working memory task performance was less coherent, in that sustained attention performance measured in the MRI session predicted later Change Localization and Square task (*ps* < 0.03, **Fig. 6A**), but not Filtering Localization, performance (*r*(18) = 0.17, *p* = 0.40, **Fig. 6A**).

We next asked if a neural signature of sustained attention measured during the fMRI session predicted subsequent long-term memory and attentional control and working memory performance. The neural model used for this purpose was the sustained attention connectome-based predictive model (saCPM; Rosenberg et al., 2016), which was constructed using fMRI connectivity. In developing this model, network nodes were defined with a 268-node functional brain atlas designed to maximize the similarity of voxel-wise time series within each node (Shen et al., 2013). For each participant, a time course was generated for each node by averaging the BOLD signal across all constituent voxels within a node at each time point during task performance. Pairwise Pearson correlation coefficients were computed between the time courses of all node pairs, and these were Fisher-normalized to produce 268 × 268 symmetric correlation matrices, reflecting each participant’s task-based functional connectivity profile. We applied the pre-trained saCPM to the functional connectivity data collected during avCPT performance from the 20 participants in our sample, namely taking the dot product of the two matrices, in generating a predicted sustained attention score for each individual.

Prior work has demonstrated that the saCPM successfully generalizes to predict sustained attention (avCPT performance) in the current dataset (Corriveau et al., 2024). In our subsample of 20 fMRI participants, saCPM-predicted sustained attention correlated with observed sustained attentional performance (*d’, r*(18) = 0.47, *p* = 0.03), validating this model as a neural proxy for sustained attention. We further investigated whether saCPM-predicted sustained attention correlated with observed measures of attentional control, working memory, and recognition memory. If sustained attention and attentional control rely on shared attentional resources in task performance, we would anticipate that saCPM predictions would be more predictive of attentional control and working memory than long-term memory performance. Conceptually replicating our findings in Experiment 1, saCPM predictions were correlated with recognition memory (*r*(18) = 0.57, *p* < 0.01, **Fig. 6B**), but not attentional control (*r*(18) = 0.15, *p* = 0.54, **Fig. 6D**), or working memory (*r*(18) = 0.28, *p* = 0.23, **Fig. 6C**) performance. Moreover, the correlation between saCPM predictions and recognition memory performance was significantly higher than the correlation between saCPM predictions and attentional control (*p* = 0.02) or working memory (*p* = 0.03) with a Fisher-Z comparison of correlation strength test. Therefore, fMRI-predicted sustained attention is more closely related to recognition memory than to attentional control and working memory.

### fMRI CPM-predicted and behaviorally measured sustained attention showed similar correlation structure with offline cognitive tasks

Lastly, we tested if CPM-predicted sustained attention resembled behavioral sustained attention within our sample. Therefore, we examined if fMRI predicted and behaviorally measured sustained attention measures shared similar covariance with attentional control, working memory, and recognition memory tasks. We calculated the Spearman correlation between the behavioral and CPM-predicted sustained attention measures and all five cognitive tasks used in our behavioral session (Change Localization and Filtering Change Localization for working memory, Flanker Square and Simon Square for attentional control, and recognition memory task). The five correlations were similarly ranked between behavioral and fMRI sustained attention measures (**Fig. 6A**). To quantify the strength of rank correlation, we found that the Spearman correlation between the five correlation coefficients using behavioral sustained attention and those using fMRI sustained attention was 0.84. Thus, fMRI-predicted sustained attention showed similar ranks in its predictive power to cognitive tasks when compared to behaviorally measured sustained attention. This suggests that the brain measures of sustained attention (i.e., CPM predictions) are specifically capturing aspects of sustained attention, as evidenced by their similar patterns of correlations with out-of-scanner task performance. In conclusion, we replicated and extended our findings in Experiment 1 that sustained attention is more closely related to long-term memory than to attentional control and working memory.

## Discussion

Here, we first established that sustained attention—although correlated with both attentional control, working memory, and long-term memory ability—forms a distinct psychological factor with a well-powered sample in Experiment 1. Contrary to an intuitive hypothesis that components of attention are more similar to each other than they are to components of memory, aggregated scores in sustained attention tasks were more predictive of long-term memory scores than attentional control and working memory scores. To test if our hypotheses could be generalized to neural patterns, we collected attentional control, working memory, and long-term memory measures 0.5-1.5 years after an independent sample of participants performed a sustained attention task during fMRI. We replicated our findings in Experiment 1—both behavioral measures of sustained attention and predictions of a validated connectome-based neuromarker of sustained attention correlated more strongly with long-term memory than attentional control and working memory measures. Therefore, we concluded that sustained attention was likely a separate factor from attention control, working memory, and long-term memory. Moreover, sustained attention differences, both behaviorally and neurally, were more predictive of long-term memory performance than attentional control and working memory task performance.

Our results suggest that attentional control and sustained attention are separable components of attention. In particular, we propose that a separate sustained attention factor, alongside a long-term memory factor and an attentional control/working memory factor, best explains the underlying factor structure of attention. The identification of the sustained attention factor supports previous claims that attentional lapses account for distinct variance from attentional control and working memory (Unsworth et al., 2021).

However, one significant difference between our studies is that the original lapse factor contained attentional lapses from a whole-report working memory task (Adam et al., 2017). Since previous research has shown that low-working-memory-capacity participants tend to experience more lapses (Adam et al., 2015), the attentional lapse factor in Unsworth et al. (2021) contains more working memory components than the factor made up of our avCPT measures. Moreover, the whole report task measured an attentional lapse by capturing trials that had one or fewer items correct. Since each trial required responses to six items in a visual array, the sampling rate for lapses was far more sparse than continuous performance tasks used in our Experiments. Therefore, future research is needed to test if continuous response tasks that capture lapse at a faster speed, and whole report tasks that measure lapses with working memory arrays over a longer time scale, require the same sustained attentional resources.

Our neural model of sustained attention did not predict offline attentional control or working memory performance as robustly as it did long-term memory measures. This finding extends previous research, which shows that brain connectome models trained with working memory abilities have significant but limited shared edges with models trained for sustained attention abilities (Avery et al., 2020; Kardan et al., 2022). Interestingly, the neural connectivity representations between working memory and fluid intelligence were significantly more similar than those with sustained attention (Avery et al., 2020). Our findings suggest that long-term memory networks may share greater variance with sustained attention than with attentional control or working memory. This result aligns with previous evidence indicating that sustained attention influences long-term memory encoding (debettencourt et al., 2018; deBettencourt et al., 2021; Wakeland-Hart et al., 2022). We speculate that the robust predictive relationship between sustained attention and long-term memory performance in our task may stem from a bidirectional interaction between these constructs. When sustained attention is high, long-term memory encoding improves. Conversely, participants with better long-term memory performance may be better able to maintain goal-relevant information, which is critical for sustained attention tasks. Further research is required to clarify the precise roles that sustained attention and long-term memory play in shaping each other’s processes.

In conclusion, we demonstrated that sustained attention, though correlated with attentional control, working memory, and long-term memory, constitutes a distinct psychological factor. Contrary to the intuitive hypothesis that attentional components are more related to each other than to memory components, sustained attention was more predictive of long-term memory than either attentional control or working memory. These findings were replicated in a follow-up study using fMRI, where both behavioral and neural markers of sustained attention correlated more strongly with long-term memory than attentional control or working memory measures. Thus, sustained attention appears to be a distinct factor that more robustly predicts long-term memory performance than attentional control and working memory. More broadly, our findings challenge conventional models that emphasize the unity of attentional processes and instead highlight the unique role of sustained attention in the formation, maintenance or even retrieval of long-term memory. These insights open new directions for research on cognitive training, educational interventions, and the neurobiological basis of sustained attention’s role in shaping memory outcomes.

## Acknowledgement

Our work was supported by Office of Naval Research Multidisciplinary University Research Initiatives (MURI) N00014-22–1-2123 to E.K.V. and M.D.R., National Science Foundation BCS-2043740 to M.D.R., and resources provided by the University of Chicago Research Computing Center.

## Appendix

**Supp. Fig 1.**
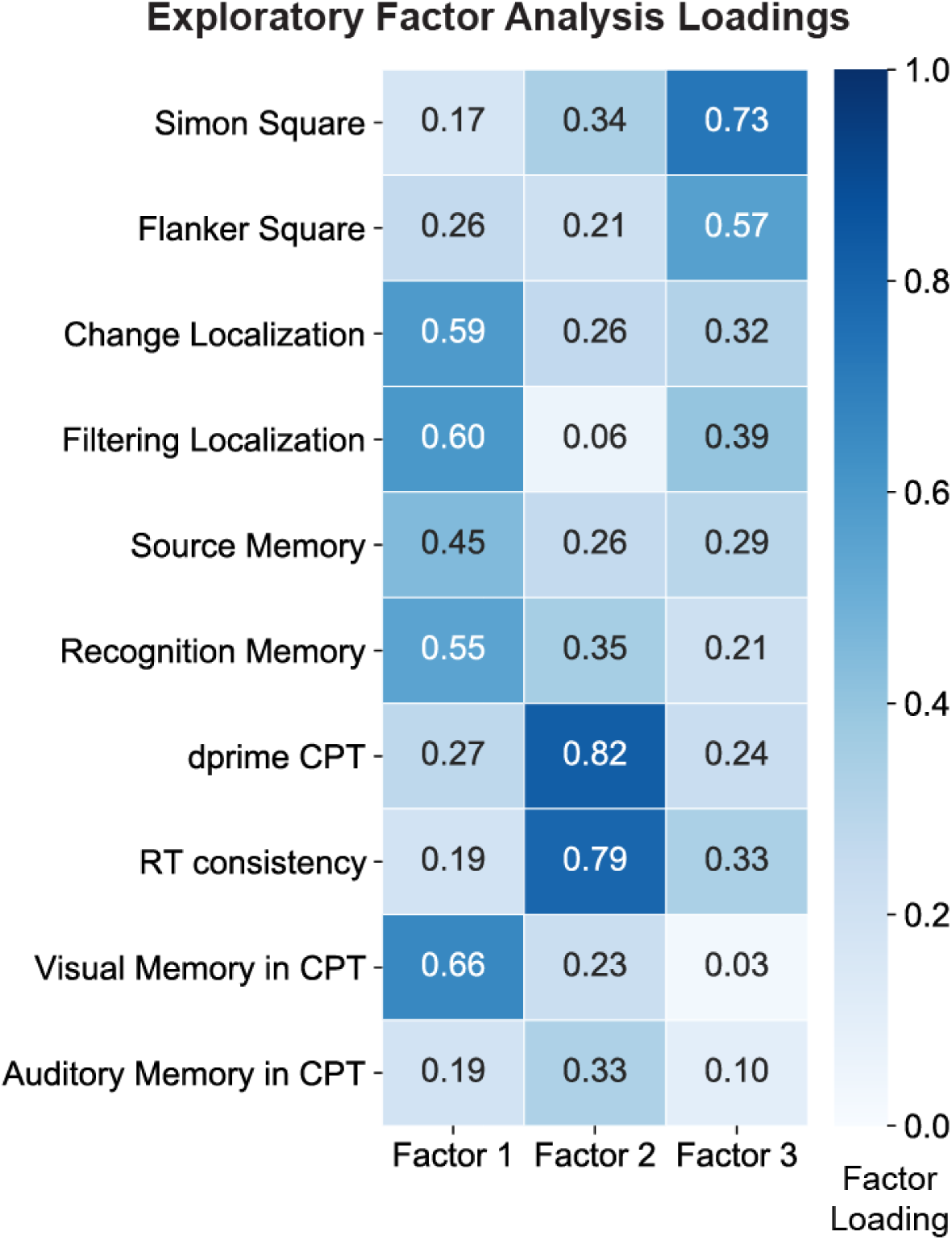
Exploratory Factor Analysis Results. The eigenvalues suggested a three-factor model. The darker the color was, the more positive the task loading was on a specific factor.

**Supp. Fig 2.**
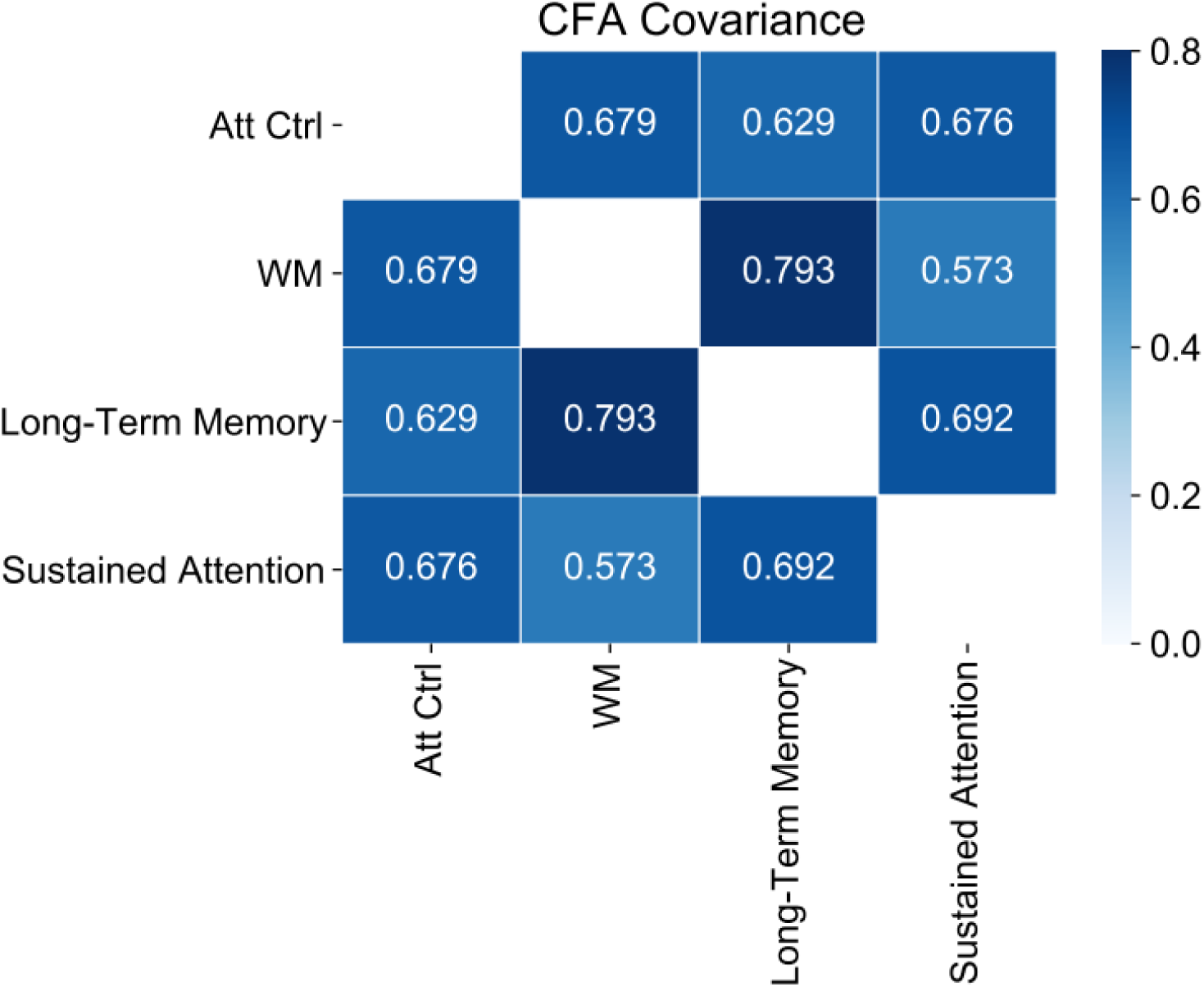
Confirmatory Factor Analysis Results with 4-factor model: Attention Control, Working Memory, Loing-Term Memory and Sustained Attention.

